# FGF Signaling Potentiates Müller Glia for Mammalian Retinal Regeneration

**DOI:** 10.64898/2026.06.26.731914

**Authors:** Neoklis Makrides, Yihua Wu, Michael Kissner, John Peregrin, Abdul Hannan, Xin Zhang

## Abstract

Müller glia possess latent regenerative potential that could be harnessed to restore retinal neurons lost to injury or disease. Although fibroblast growth factor (FGF) signaling is upregulated following retinal damage, its role in mammalian retinal regeneration remains unclear. Here, we investigated the function of FGF signaling in Müller glial reprogramming using genetic, pharmacological, and single-cell transcriptomic approaches. Activation of FGF signaling alone was insufficient to induce Müller glial proliferation in the mouse retina. However, conditional deletion of FGFR1/2 in Müller glia abolished regenerative responses induced by multiple independent pathways, demonstrating that FGF signaling is essential for regenerative competence. Mechanistically, loss of FGF signaling impaired sustained ERK/MAPK activation following injury, while MEK/ERK inhibition phenocopied the regenerative defect. Conversely, constitutive MEK activation induced limited Müller glial proliferation in the absence of injury. Although STAT3/5 inhibition synergized with Activin-A to promote robust proliferation and neurogenic gene expression, it failed to rescue regeneration in FGF-deficient Müller glia. Single-cell RNA sequencing revealed that FGF signaling suppresses multiple anti-regenerative programs, including Hes1, S1pr1, p27, NF-κB, and p300/CBP activity. Together, these findings identify FGF signaling as a critical permissive regulator of mammalian retinal regeneration that potentiates Müller glia through sustained ERK activation and suppression of transcriptional barriers to regeneration.

## Introduction

A major goal of vision restoration research is to stimulate retinal Müller glia to proliferate and differentiate into the neuronal populations lost during degenerative disease or injury. The facultative stem cell properties of Müller glia were first described in lower vertebrates, including teleost fish and chick, where these cells can spontaneously regenerate the retina following damage ^1–6^. Although this regenerative capacity has largely been lost during mammalian evolution, reactivation of signaling pathways that govern regeneration in zebrafish and chick has been shown to induce limited regenerative responses in mammals. Several pathways have been implicated in this process, including Wingless-related integration site (Wnt) ^7–10^, Epidermal Growth Factor (EGF) ^11–17^, Hippo/YAP ^18^, and Notch signaling ^10^. Furthermore, forced expression of neurogenic transcription factors such as *Ascl1* ^19,20^ and *Neurod1* ^21,22^ can reprogram mammalian Müller glia toward a regenerative state. An important distinction between mammalian and non-mammalian regeneration is that Müller glial proliferation in mammals generally occurs only following retinal injury, whereas in zebrafish, exogenous administration of growth factors such as EGF and Ciliary Neurotrophic Factor (CNTF) is sufficient to induce Müller glial dedifferentiation in the absence of injury ^23,24^. This observation has led to the hypothesis that mammalian Müller glia require both growth factor signaling and injury-associated inflammatory cues to become regeneration-competent. Among the pathways activated following retinal injury, Fibroblast Growth Factor (FGF) signaling is consistently upregulated across multiple species, suggesting a potentially conserved role in retinal repair and regeneration ^25–29^.

The FGF family was originally identified from bovine pituitary extracts based on its ability to stimulate fibroblast proliferation in culture ^30^. The family comprises 18 ligands grouped into six subfamilies, each exhibiting distinct affinities for the four FGF receptors ^31–33^. Given their potent mitogenic activity and essential role during retinal development ^34^, FGFs have long been considered attractive candidates for promoting retinal regeneration. Indeed, their proliferative effects have been demonstrated in vitro, where treatment with basic FGF (bFGF) markedly increases proliferation of bovine- and human-derived Müller glia ^35–37^.

Despite the conserved mitogenic activity of FGF signaling in vitro, its role in retinal regeneration in vivo remains controversial. In zebrafish, intravitreal administration of bFGF following light-induced retinal injury significantly enhanced Müller glial regeneration ^38^. Furthermore, combined treatment with bFGF and interleukin-6 induced Müller glial proliferation in goldfish even in the absence of injury ^39^. Conversely, genetic overexpression of FGF8 in zebrafish reduced Müller glial proliferation after injury ^40^. Subsequent analyses revealed that this inhibitory effect was age-dependent, as forced FGF8 expression promoted proliferation in younger fish ^40^. Despite these context-dependent outcomes, loss-of-function studies consistently demonstrated that pharmacological inhibition of FGF signaling following injury reduced regenerative efficiency in adult zebrafish ^38,41^. Collectively, these findings indicate that FGF signaling contributes to retinal regeneration in teleosts, although its precise role appears highly dependent on developmental stage and cellular context.

Studies in the chick retina similarly support a role for FGF signaling during Müller glia–mediated regeneration. Intraocular administration of bFGF following excitotoxic injury induced modest Müller glial proliferation ^42,43^, which was substantially enhanced when combined with insulin, insulin-like growth factor-1 (IGF-1) ^28,42,44,45^, the retinoic acid agonist TTNPB ^46^, or the Smad3 inhibitor SIS3 ^43^. Conversely, inhibition of BMP or Notch signaling using DMH1 or DAPT reduced the regenerative effects of bFGF ^43,45^. These observations suggest that while FGF acts as a mitogenic stimulus, additional signaling pathways regulate Müller glial dedifferentiation and regenerative competence. Supporting this notion, retinoic acid activation combined with bFGF induced robust Müller glial proliferation in the absence of retinal injury ^46^. However, studies involving microglia-depleted chick retinas have complicated this interpretation. In these models, spontaneous regeneration following injury was impaired, indicating that Müller glial priming depends on signals derived from activated microglia ^47^. Notably, administration of either heparin-binding EGF (HB-EGF) or FGF restored regenerative capacity in microglia-depleted retinas, suggesting that FGF may function as a permissive or priming factor rather than solely as a mitogen ^47^. Consequently, it remains unclear whether FGF primarily promotes proliferation, potentiates regenerative competence, or performs both functions during retinal regeneration. Importantly, both fish and chick retain an intrinsic capacity for spontaneous retinal regeneration, raising the possibility that Müller glial dedifferentiation in these species occurs independently of many injury-associated signals that are required in mammals.

Compared with fish and chick, relatively little is known about the role of FGF signaling during mammalian retinal regeneration. Nevertheless, co-administration of bFGF and insulin following retinal injury in mice induces only modest Müller glial proliferation and fails to generate substantial neuronal differentiation ^13,48^. Interestingly, the neurogenic potential of this treatment is significantly enhanced in mice lacking Nfia, Nfib, and Nfix, suggesting that FGF primarily exerts a proliferative rather than neurogenic effect in mammalian Müller glia ^49^. Given the conflicting findings across species and the incomplete understanding of FGF function during retinal regeneration, we sought to define both the necessity and sufficiency of FGF signaling in promoting Müller glial regeneration in the mouse retina and to identify the downstream molecular pathways involved. We demonstrate that FGF functions as an essential niche-derived signal that potentiates Müller glial dedifferentiation by suppressing STAT3 activity and p300/CBP-dependent transcriptional programs. Furthermore, combined Activin-A stimulation and STAT3/5 inhibition produced the most robust Müller glial proliferation observed in our study. However, this treatment failed to rescue regeneration in FGF-deficient Müller glia, indicating that FGF-mediated potentiation extends beyond STAT-dependent mechanisms. Together, these findings establish FGF signaling as a critical regulator of regenerative competence in the mammalian retina.

## Materials and methods

### Mice husbandry

All husbandry and experimental procedures performed on the mice were approved by the Columbia University’s Institutional Animal Care and Use Committee and conform to the relevant regulatory standards. The mice used for this study were *Rax-CreER^T2^* (obtained from Seth Blackshaw, RRID:IMSR_JAX:025521), *Fgfr1^flox^* (Jackson Laboratory, RRID:MGI:3817869), *Fgfr2^flox^* (obtained from Dr. David Ornitz, Washington University at St. Louis), *R26*^LSL-Fgf8^ (obtained from Liang Ma, Washington University at St. Louis), *Ptpn11^flox^* ^50^, *R26^LSL-MEK1DD^* (Jackson Laboratory, Stock No: 012352) ^51^ and *Ai9* (Jackson Laboratory, RRID:MGI:3817869). All animals were kept in a humidity-controlled environment at a constant temperature of 21°C and were subjected to a 12:12 light/dark cycle and with access to water and conventional chow diet ad libitum. The mice used for the study were kept in a C57BL/6 background and were initially screened for the common retinal degeneration mutations. Only male mice were included in the experimental procedures and cre negative littermates were used as controls. The genotype of the mice was determined through standard PCR using the primers listed on Supplementary Table S1.

### Intraocular Injections

Adult mice were anesthetized by intraperitoneal injection of ketamine/xylazine (071069 and 061035, Covetrus) and sterile tropicamide (070498, Covetrus) and phenylephrine hydrochloride (068882, Covetrus) drops were applied to the eye, followed by proparacaine hydrochloride (068926, Covetrus) for topical anaesthesia. Lubricant eye gel was applied on each eye to prevent corneal ulcers and using a sterile acupuncture needle, a small hole was created at the corneoscleral boundary. A glass capillary needle was used to inject 1.5 μl of each treatment solution at a concentration indicated on Table S2. Treatment injections were only performed on the right eye of mice while the left eye was used as the control solvent injection. Proparacaine hydrochloride eye drops and lubricant eye gel were applied again on each eye and the mice were placed on a heating pad to recover.

### Tissue harvest and Immunohistochemistry

The mice were euthanised through cervical dislocation and the eyes were enucleated and fixed in 4% paraformaldehyde (PFA) overnight at 4°C. The eyes were washed three times in Phosphate Buffer Solution (PBS) and the cornea and lens were subsequently removed. Eyes that were used for cryosectioning, were cryoprotected in 30% Sucrose solution (in PBS) overnight at 4°C and thereafter embedded in cryomolds containing Tissue-Tek® optimal cutting temperature (OCT) compound (Sakura, Japan). The tissues were serially sectioned at 10 μm thickness in a transverse plane using a Leica CM1850 UV-3-1 Cryostat Microtome (Leica Biosystems, US). Eyes used for flat mount staining were cut in four quadrants (superior, inferior, nasal and temporal) and the retina was separated from the Retinal Pigmented Epithelium (RPE). The flat mounts were permeabilized in 0.03% triton-X (in PBS) before immunostaining. Both retinal flat mounts and rehydrated cryosections were blocked with 10% horse serum for 1 h at room temperature and incubated with a primary antibody (diluted in blocking solution) overnight at 4°C. The tissues were washed three times in PBS for 5 min and were incubated with the secondary fluorophore-conjugated antibody in blocking solution for 1 h at room temperature in the dark. The Samples were washed and mounted with n-propyl gallate anti-fading reagent (P3130, Sigma-Aldrich) and examined under a Leica DM5000-B fluorescence microscope. The antibodies used for the study are listed on Table S3.

### EdU incorporation assay & TUNEL staining

Mice were injected with EdU (ab146186, Abcam) dissolved in DMSO at the dosage of 50 mg/kg body weight for two consecutive days following the intraocular injections. For EdU detection, the Click-IT EdU Imaging Kit (C10337, Invitrogen) was used according to the manufacturer’s instructions. TUNEL staining was performed on cryosections using the Apo-Direct TUNEL assay kit according to the manufacturer’s instructions (APT110, Sigma-Aldrich).

### Western blot

Eyes were enucleated as previously described and the cornea and lens were dissected out in ice cold PBS. The retina was removed from the RPE and lysed in in CelLytic-M lysis buffer (Sigma) with proteinase inhibitor cocktail (Thermo fisher). The lysates were sonicated and centrifuged at 12,000 g for 10 min. The supernatant was collected and loading buffer with the anionic detergent sodium dodecyl sulfate (SDS) was added to the samples. The mixture was boiled at 95°C for 5 min and equal amounts of total protein were run on an SDS-Page gel. The membranes were subsequently stained according to standard protocols and visualized using an Odyssey SA scanner (LICOR Biosciences, Lincoln, NE).

### Electroretinography (ERG)

ERG recordings were performed using the Celeris rodent ERG system (Diagnosys, Lowell, MA). The mice were dark adapted for at least 12 h before the procedure. To maintain a constant body temperature during the procedure, the animals were kept on a heating pad (Mycoal, Tochigi, Japan). Mice were anesthetized and prepped as previously described and initially subjected to a scotopic 0.9 log cd.s/m^2^ stimulus repeated 10 times. Following the completion of the dark-adapted recordings, the animals were exposed to a full-field 30 cd/m^2^ white background for 10 min. The photopic recordings were performed along this continuous background. A total of 10 repeats of 1.47 log cd.s/m^2^ stimuli were performed and a band-pass filter ranging from 0.3 to 300 Hz was used to isolate the signals from the recorded waves.

### Optical Coherence Tomography (OCT) imaging

OCT imaging was performed using the OCT-SLO Spectralis 2 scanning laser confocal ophthalmoscope (Heidelberg Engineering, Heidelberg, Germany). Mice were anesthetized and prepped as previously described and a heating pad was used to maintain constant body temperature during the procedure. The mice were screened using a 55° widefield objective lens with a working distance of 13.5 mm.

### Single Cell RNA sequencing

Single-cell sequencing was performed at the single-cell sequencing core in Columbia Genome Center. Whole retinas were dissociated using 40 U/ml papain (LS003119, Worthington Biochemical, Lakewood, NJ) according to previous protocols ^52^. The dissociated cells were loaded into a chromium microfluidic chip with v3 chemistry and barcoded with a 10× chromium controller (10× Genomics). The samples were reverse-transcribed and the Chromium Single Cell v3 reagent kit was used to generate the sequencing libraries (10× Genomics) which were subsequently sequenced on the NovaSeq 6000 (Illumina). The gene-cell matrices were imported in R and analysed using the Seurat toolkit. Low-quality cells were filtered out (<200 or > 4000 transcripts/cell, and >25% mitochondria genes) and the gene expression levels was normalized using the “NormalizedData” function. Doublets were excluded using the ‘DoubletFinder’ R package (v2.0.3). Cell clustering was visualised though Uniform manifold approximation and projection (UMAP) and heatmaps were created using the “doHeatmap” function. RNA velocity analysis was carried out in pythons using the velocyto (v0.17.17) and scvelo (v0.0.4) packages. Regulon analysis was performed using the pySCENIC (v1.1.3) packages in python. The RNAseq data were deposited at the Gene Expression Omnibus (GEO) database.

### Statistical analysis

The quantification of the GFAP relative expression area was performed using ImageJ. The mean values obtained from four sections within the same animal were considered as a single biological replicate. A total of three biological replicates were analysed to assess statistical differences. The regeneration efficiency of each treatment was evaluated by counting the total number of Sox2/Edu^+^ and Sox2^+^ cells in either whole retinal flat mounts or from the mean value of every fifth section of an entire eye (that correspond to 40-50 sections) accordingly. The ratio of the Sox2/Edu^+^ cells with the total number of Sox2 cells was used to distinguish differences between wild type and conditional knockout retinas by the Mann-Whitney U Test due to the small sample size of each group. To test differences between multiple treatments, Kruskal-Wallis one-way ANOVA was used. Retinas that exhibited extensive damage or retinal detachment were excluded from the study. Signal intensity from western blot images of the phosphorylated protein and total protein was quantified using ImageJ. The Ratios between the phosphorylated and unphosphorylated protein levels were used to test differences between wildtype and conditional knockout mice using Mann-Whitney U Test.

## Results

### FGF Signaling Alone Is Insufficient to Promote Müller Glia Regeneration in Adult Mice

To determine whether FGF signaling is sufficient to induce retinal regeneration, adult mice (P60) received intravitreal injections of basic FGF (bFGF) for two consecutive days in the absence of excitotoxic injury. Concurrently, animals were administered EdU intraperitoneally to assess cellular proliferation. A short EdU-labelling period was selected to minimize cumulative incorporation bias and provide a more accurate assessment of proliferative activity. To confirm activation of the FGF pathway, we first examined the downstream ERK signaling cascade. Basal levels of phosphorylated ERK (p-ERK) were low in Müller glia of untreated retinas; however, bFGF administration induced robust p-ERK expression that colocalized with Sox2-positive Müller glia (Fig. 1A). These findings confirmed that intravitreal bFGF delivery effectively activated FGF signaling in retinal Müller glial cells. Despite efficient pathway activation, FGF stimulation did not induce Müller glial proliferation. Although a small number of EdU-positive cells were detected throughout the retina (green arrows), none co-expressed the Müller glial marker Sox2 (Fig. 1A). Instead, these EdU-positive cells colocalized with IBA1, indicating that they were proliferating microglia rather than Müller glia (Fig. S1A). To further validate this observation while eliminating potential confounding effects associated with intravitreal injections, we conditionally overexpressed FGF8 specifically in Müller glia using the *Rax-CreER^T2^* mouse line. Consistent with the bFGF injection experiments, no Sox2+/EdU+ cells were detected following tamoxifen-induced FGF8 expression (Fig. 1B). Although FGF8 induction resulted in a modest increase in p-ERK levels, the magnitude of activation was lower than that observed following exogenous bFGF administration (Fig. 1C). Thus, activation of FGF signaling alone was insufficient to stimulate Müller glial cell-cycle re-entry.

**Figure 1.**
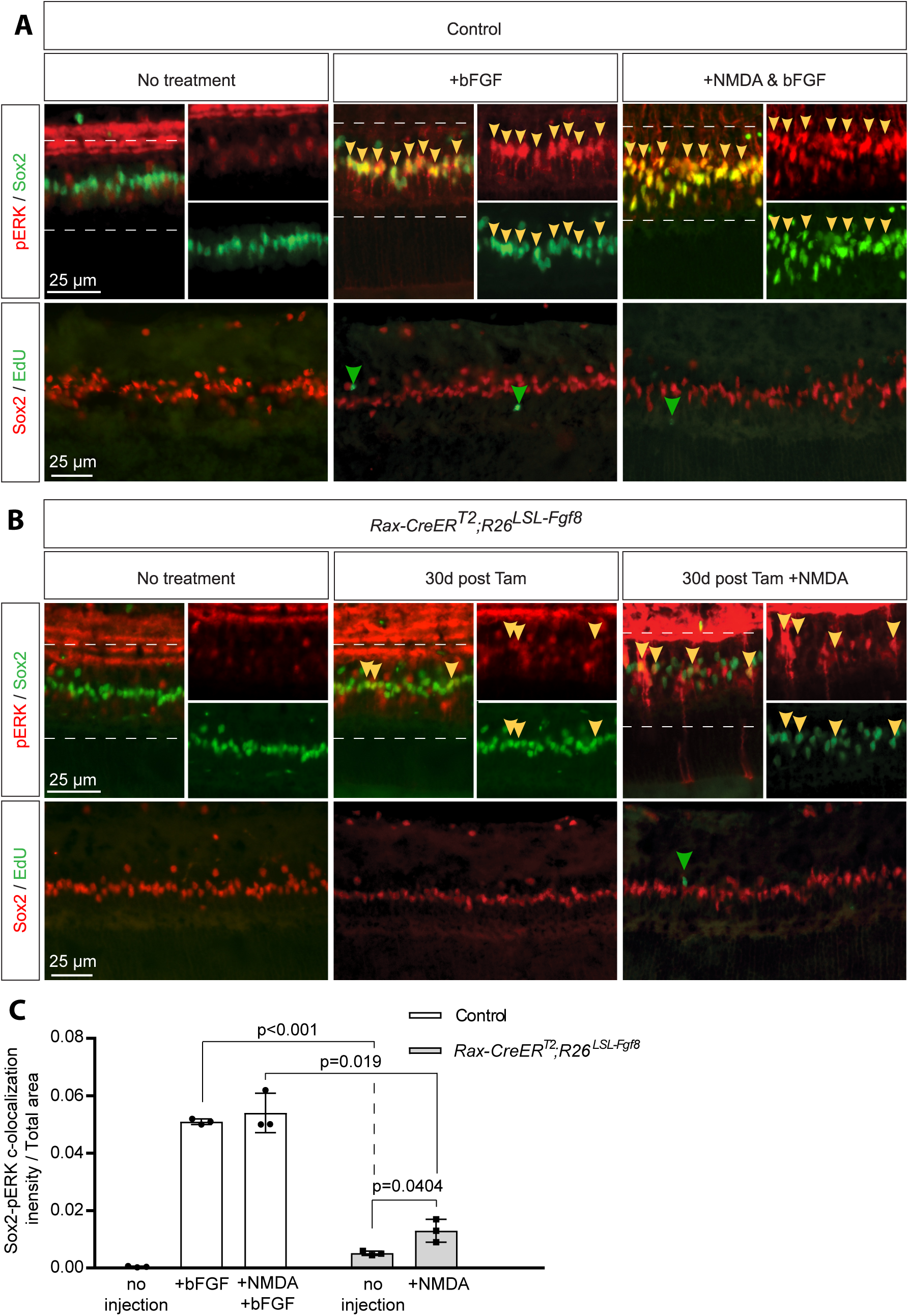
FGF signalling alone is insufficient to induce Müller glial proliferation. **(A)** Immunostaining for Sox2 and phosphorylated ERK (pERK) in adult mouse retinas under basal conditions or 30 min after intravitreal bFGF injection. **(B)** Immunostaining for Sox2 and EdU in control and *Rax-CreER^T^*^2^;*R26^LSL-Fgf8^*retinas 30 days after tamoxifen induction, with or without NMDA injury. **(C)** Quantification of pERK-positive Müller glia shown as Sox2/pERK colocalization (mean ± S.D., One-way ANOVA, *n* = 3 biological replicates).

We next investigated whether enhanced FGF signaling could promote regeneration in the context of acute retinal injury. Prior to these experiments, the efficacy of the excitotoxic injury model was confirmed, as a single intravitreal injection of NMDA resulted in substantial depletion of retinal ganglion cells, evidenced by the loss of Brn3a immunoreactivity (Fig. S1B). However, neither exogenous bFGF administration nor Müller glia-specific FGF8 overexpression following NMDA-induced injury increased Müller glial proliferation, as no Sox2+/EdU+ cells were detected under either condition (Fig. 1A,B). Collectively, these findings demonstrate that although FGF signaling effectively activates downstream mitogenic pathways in Müller glia, it is not sufficient to induce Müller glial proliferation or regeneration in the adult mouse retina, either in the absence or presence of acute retinal injury.

Previous studies in zebrafish have suggested that the regenerative effects of FGF signaling are age dependent. We therefore examined whether younger mammalian retinas exhibit greater responsiveness to FGF stimulation. To test this possibility, bFGF was administered intravitreally to postnatal day 16 (P16) and postnatal day 21 (P21) mice. Similar to the results observed in adult animals, no Sox2+/EdU+ Müller glia were detected following FGF treatment at either developmental stage (Fig. S1C). These data indicate that, unlike in certain teleost models, the inability of FGF signaling to induce Müller glial proliferation is not overcome in younger mouse retinas.

### Ablation of FGF Signaling in Müller Glia Induces a Gliotic Response

Given that FGF signaling alone was insufficient to stimulate retinal regeneration, we next investigated whether it is required for this process. To address this question, we generated a Müller glia-specific conditional knockout of *FGFR1* and *FGFR2*, the two dominant FGFR in the Müller glia (Fig. S1D). Before examining the regenerative consequences of FGF receptor deletion, we first validated the efficiency and specificity of Cre-mediated recombination using the Ai9 reporter line. Tamoxifen administration at P30 resulted in specific labelling of more than 90% of Müller glia, as demonstrated by the extensive colocalization of tdTomato with the Müller glial markers Sox9, Sox2, and CyclinD3 (Fig. 2A,B). This high degree of specificity confirmed the suitability of the *Rax-CreER^T2^* driver for subsequent lineage-tracing and conditional knockout experiments. Importantly, deletion of *FGFR1/2* did not alter the number of tdTomato-positive Müller glia, indicating that FGF signaling is dispensable for Müller glial survival under homeostatic conditions (Fig. 2A,B). Efficient disruption of FGF signaling was confirmed by western blot analysis of retinal lysates collected 30 minutes after intravitreal bFGF administration. Conditional deletion of *FGFR1/2* markedly reduced ERK activation in response to bFGF stimulation, demonstrating successful inhibition of downstream FGF signaling (Fig. 2C,D).

**Figure 2.**
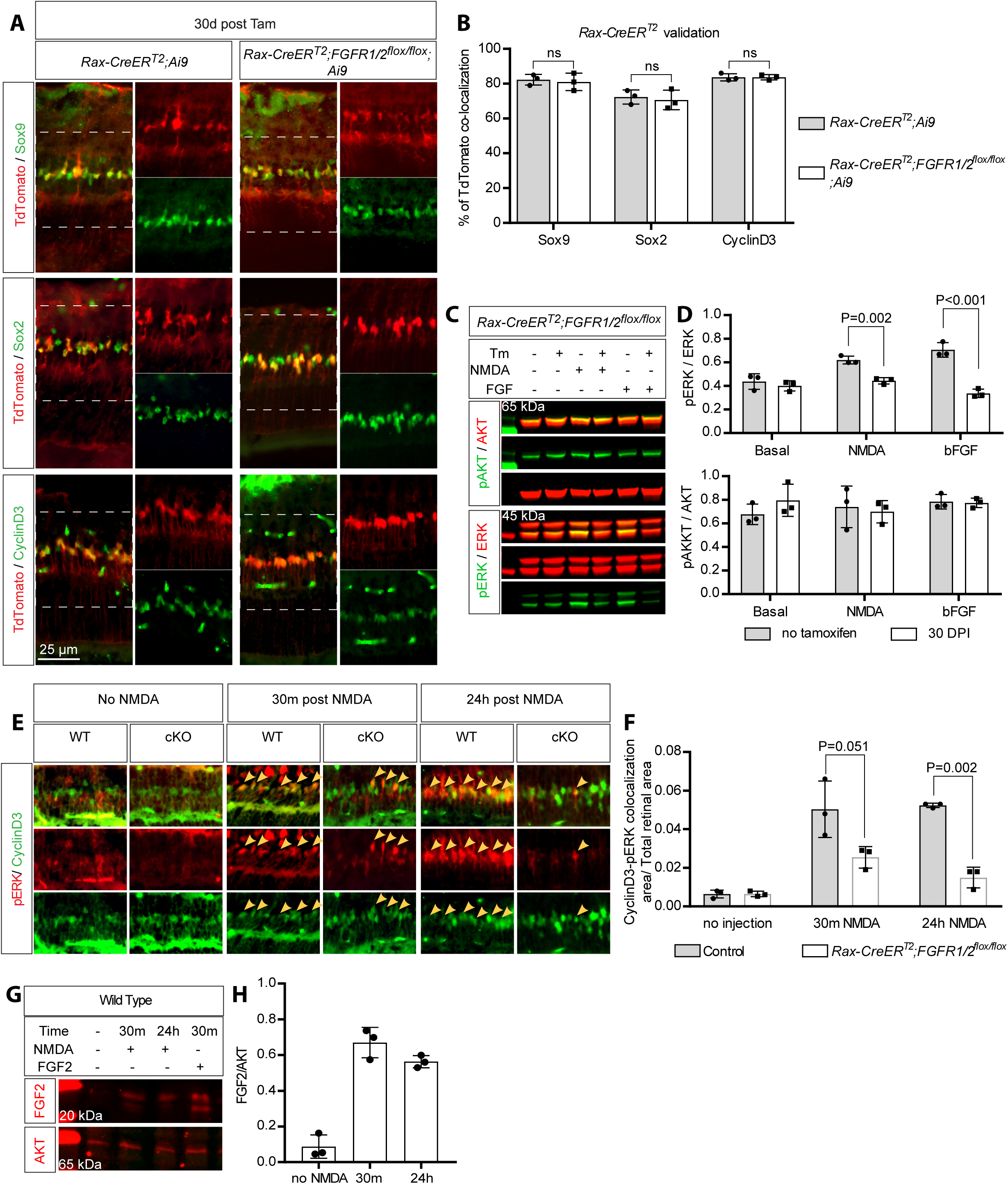
Conditional deletion of FGFR1/2 abolishes injury-induced ERK activation in Müller glia. **(A)** Validation of *Rax-CreER^T2^* recombination specificity using the *Ai9* reporter in adult control and *FGFR1/2* conditional knockout retinas stained for CyclinD3, Sox9 and Sox2. **(B)** Percentage of tdTomato-positive cells co-expressing Müller glial markers (mean ± S.D., Two-sided Student’s t test, ns: not significant, *n* = 3). **(C)** Representative immunoblots of pERK and pAKT from control and *Rax-creER^T2^;Fgfr1^flox/flox^;Fgfr2^flox/flox^*retinal lysates collected 30 min after NMDA or bFGF injection. **(D)** Quantification of phosphorylated protein levels relative to total protein (mean ± S.D., Two -sided Student’s t test, *n* = 3). **(E)** Immunostaining for pERK and CyclinD3 at 30 min and 24 h following NMDA injury (yellow arrows indicate colocalization). **(F)** Quantification of pERK-positive Müller glia (mean ± S.D., Two -sided Student’s t test, *n* = 3). **(G)** Representative immunoblot showing FGF2 expression following NMDA injury; bFGF-treated retinas served as positive controls. **(H)** Quantification of FGF2 protein levels normalized to total AKT (mean ± S.D., Two -sided Student’s t test, *n* = 3).

Interestingly, reduced ERK phosphorylation was also observed in conditional knockout retinas following NMDA-induced injury alone (Fig. 2C,D). Under normal conditions, p-ERK expression was largely restricted to the plexiform layers; however, acute retinal injury induced a rapid and pronounced increase in p-ERK within Müller glia (Fig. 2E,F). Elevated ERK activation persisted for at least 24 hours following injury in control retinas. In contrast, Müller glia lacking *FGFR1/2* exhibited substantially reduced ERK activation shortly after NMDA administration, and only limited p-ERK immunoreactivity remained detectable after 24 hours. These findings suggest that FGF signaling represents a major receptor tyrosine kinase pathway activated in Müller glia following retinal injury. Consistent with this interpretation, western blot analysis revealed a significant increase in FGF2 protein levels after NMDA-induced damage (Fig. 2G,H).

We next examined the consequences of FGF signaling ablation on retinal homeostasis. Unexpectedly, conditional deletion of *FGFR1/2* resulted in a progressive gliotic response, characterized by a gradual increase in GFAP expression within Müller glia (Fig. 3A). Quantification revealed that this gliotic response reached a plateau approximately three weeks after tamoxifen induction (Fig. 3B). Furthermore, following NMDA-induced injury, GFAP upregulation was markedly more pronounced in conditional knockout retinas than in wild-type controls (Fig. 3C,D), suggesting that loss of FGF signaling sensitizes Müller glia to injury, potentially due to an underlying state of cellular stress.

**Figure 3.**
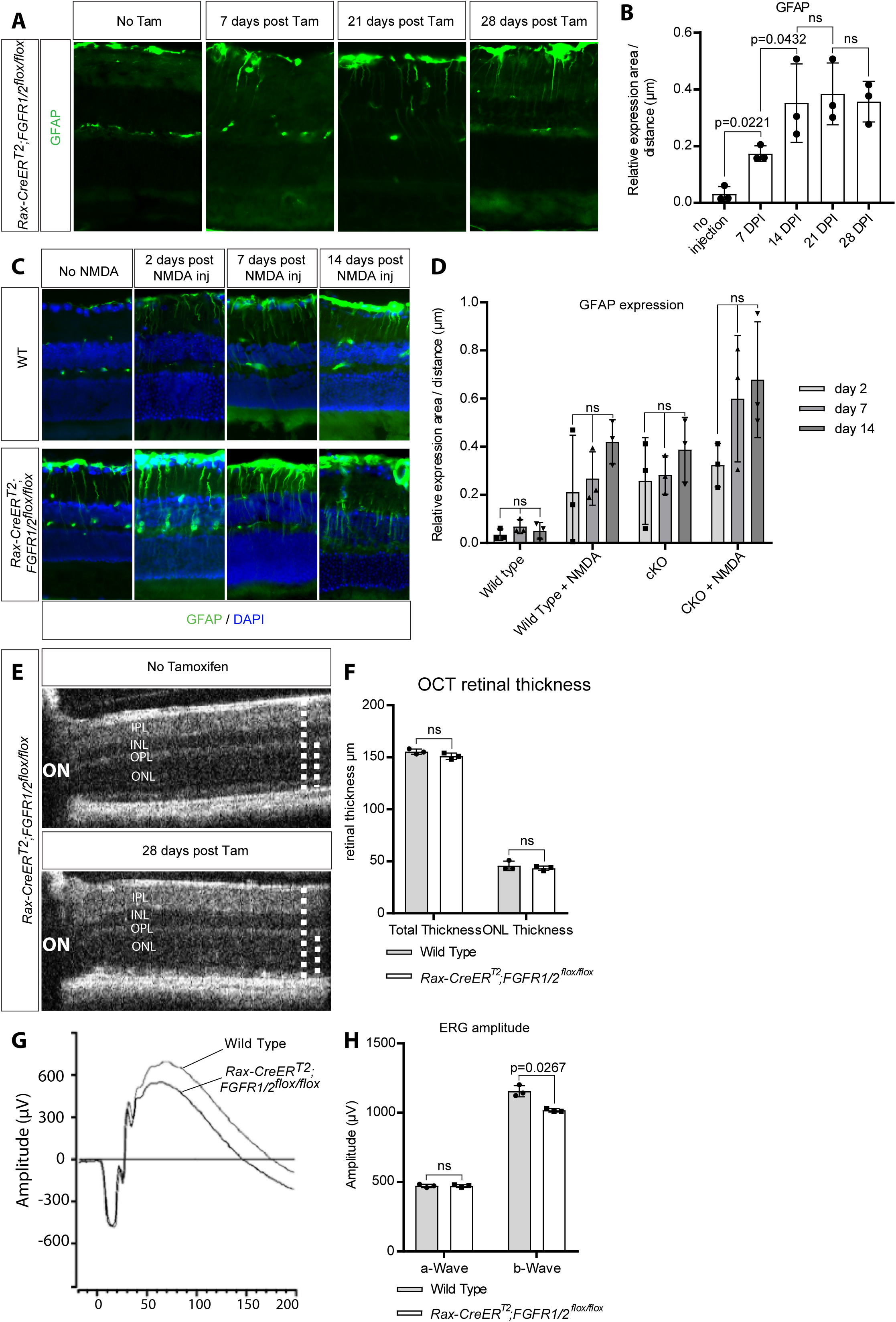
Loss of FGF signalling induces reactive gliosis in Müller glia. **(A)** GFAP immunostaining showing progressive Müller glial gliosis following conditional deletion of *FGFR1/2*. **(B)** Quantification of GFAP fluorescence intensity (mean ± S.D., One-way ANOVA, ns: not significant, *n* = 3). **(C)** GFAP immunostaining following NMDA-induced injury in control and *FGFR1/2* conditional knockout retinas. **(D)** Quantification of injury-induced GFAP expression (mean ± S.D., One-way ANOVA, ns: not significant, *n* = 3). **(E)** Representative OCT images of control and mutant retinas. **(F)** Quantification of retinal thickness measured by OCT (mean ± S.D., Two -sided Student’s t test, ns: not significant, *n* = 3). **(G)** Representative electroretinogram (ERG) recordings. **(H)** Quantification of ERG amplitudes (mean ± S.D., Two -sided Student’s t test, ns: not significant, *n* = 3).

Despite the development of reactive gliosis, the structural integrity of the retina remained largely preserved. Optical coherence tomography (OCT) analysis revealed no significant changes in retinal thickness up to four weeks following *FGFR1/2* deletion (Fig. 3E,F). Similarly, electroretinography (ERG) recordings showed only a modest reduction in retinal response amplitudes in mutant mice compared with controls at this time point (Fig. 3G,H). These functional deficits likely reflect the effects of Müller glial gliosis rather than overt retinal degeneration. Collectively, these results demonstrate that FGF signaling is a major injury-responsive pathway in Müller glia and plays an important role in maintaining their homeostatic state by suppressing reactive gliosis.

### FGF Signaling Is Required for Müller Glia Regeneration Through Sustained ERK Activation

Having established that conditional ablation of FGF signaling did not affect Müller glial survival, we next examined whether FGF signaling is required for Müller glia regeneration. To address this question, we employed several independent regeneration paradigms to determine whether FGF signaling functions as a common mediator of regenerative responses induced by distinct signaling pathways. Similar to the previous FGF stimulation experiments, regenerative factors were administered by intravitreal injection on two consecutive days beginning two days after NMDA-induced injury. EdU was administered concurrently over the same period to assess proliferative activity while minimizing cumulative labelling bias. Consistent with previous reports, administration of EGF, CHIR99021, or SB431542 induced Müller glial proliferation, as demonstrated by the presence of Sox2+/EdU+ cells within the retina (Fig. 4A,B). In addition, intravitreal delivery of Activin-A or the ROCK1/2 inhibitor Y-27632 similarly promoted Müller glial cell-cycle re-entry. Among all treatments tested, Activin-A produced the greatest proliferative response, although this increase did not reach statistical significance relative to the other regeneration paradigms.

**Figure 4.**
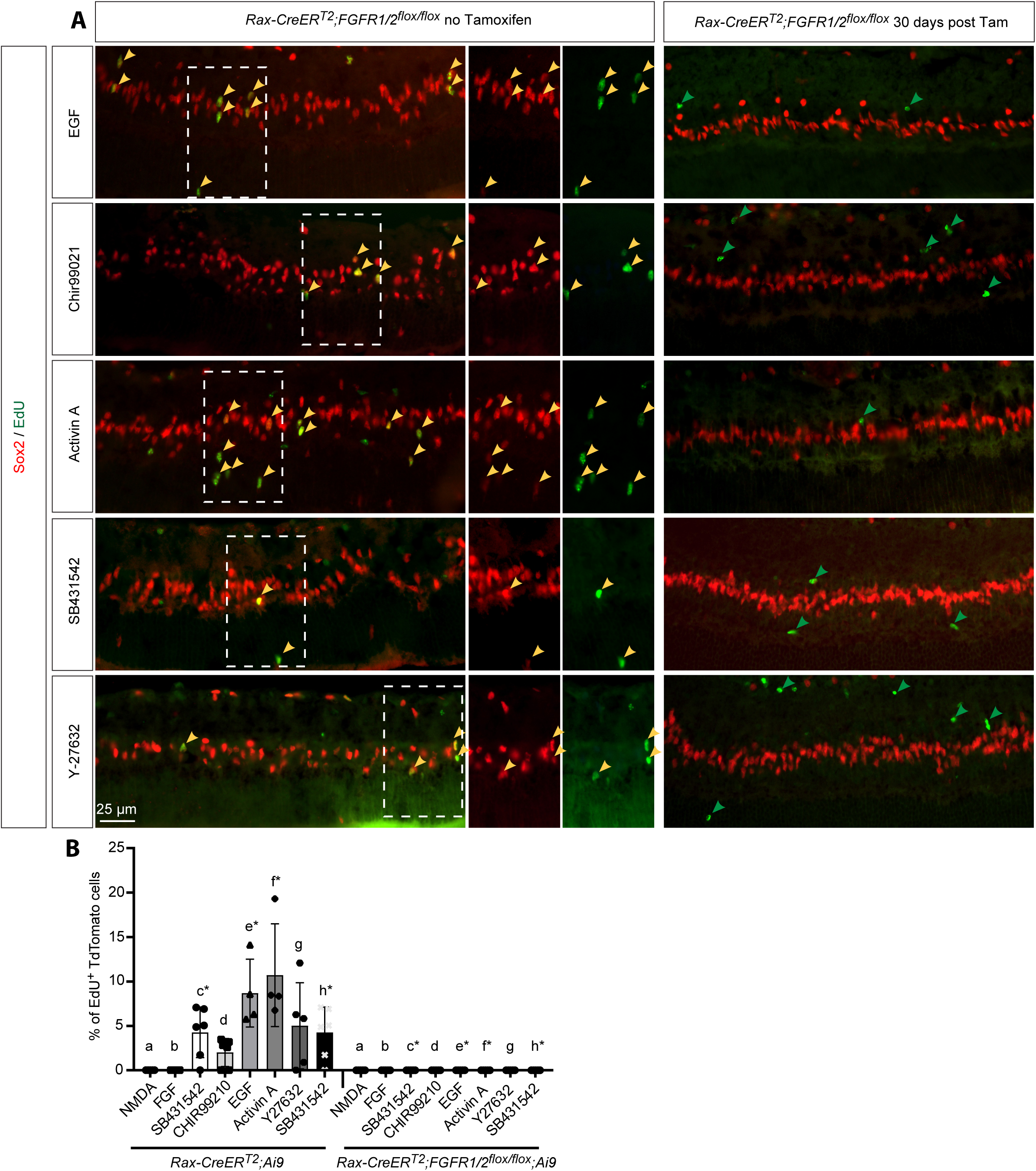
FGF signalling is required for Müller glial regeneration. **(A)** Representative retinal sections stained for Sox2 and EdU following treatment with EGF, CHIR99021, Activin-A, SB431542 or Y-27632 after NMDA injury in control and FGFR1/2 conditional knockout mice. **(B)** Quantification of proliferating Müller glia shown as Sox2+/EdU+ cells (mean ± S.D., One-way ANOVA, asterisk denotes significant difference between control and comparable treated conditional knockout, *n* = 4–5 biological replicates).

We next investigated whether these regenerative responses were dependent on FGF signaling. To minimize potential confounding effects arising from the progressive gliosis observed following receptor deletion, tamoxifen was administered three weeks before NMDA injury, allowing sufficient time for efficient FGFR ablation while maintaining retinal integrity. Strikingly, although EdU-positive and Sox2-negative microglia remained readily detectable in the conditional knockout retinas (Fig. 4A, green arrows), proliferating Müller glia were completely absent across all regenerative treatments examined, including EGF, Activin-A, and CHIR99021 (Fig. 4A,B). Furthermore, FGFR-deficient Müller glia exhibited impaired nuclear migration compared with their wild-type counterparts (Fig. 4A), suggesting a broader disruption of the regenerative response.

The two principal downstream effectors of FGF signalling are the RAS/MAPK and PI3K/AKT pathways. Consistent with efficient receptor ablation, activation of both ERK and AKT was markedly reduced in FGFR-deficient Müller glia under all regenerative conditions examined, whereas induction of the Wnt target Lef1 following CHIR99021 treatment remained unaffected (Fig.S2A-B”). To further determine the contribution of the RAS/MAPK and PI3K/AKT pathways to Müller glial regeneration, we generated Müller glia-specific Shp2 conditional knockout mice (*Rax-CreER^T2^*;*PTPN11^flox/flox^*), in which activation of both pathways downstream of receptor tyrosine kinases is disrupted. Tamoxifen was administered three weeks prior to retinal injury and regenerative stimulation. Similar to the FGFR knockout model, Shp2-deficient Müller glia failed to proliferate in response to EGF, CHIR99021, or Activin-A treatment (Fig. S3A). Consistent with successful pathway disruption, activation of both the PI3K/AKT and MEK/ERK cascades was markedly reduced in these retinas (Fig. S3A). To distinguish the relative contributions of each pathway, regenerative stimuli were co-administered with either the PI3K inhibitor LY294002 or the MEK inhibitor PD0325901 in wild-type mice (Fig. S3B-E). Pharmacological inhibition of PI3K significantly reduced, but did not completely abolish, Müller glial proliferation induced by EGF, CHIR99021, or Activin-A (Fig. S3B,C). In contrast, inhibition of MEK/ERK signaling completely prevented regeneration, with no Sox2+/EdU+ Müller glia detected under any treatment condition (Fig. S3B,C). Notably, Müller glial nuclear migration was still observed following inhibition of either pathway despite the marked reduction in proliferative activity (Fig. S3B), indicating that nuclear translocation and cell-cycle entry are regulated by distinct molecular mechanisms.

Together, these findings demonstrate that sustained MEK/ERK activation is essential for Müller glial proliferation during retinal regeneration, whereas PI3K/AKT signaling plays a supportive but non-essential role. To further test whether prolonged ERK activation is sufficient to promote regenerative competence, we constitutively activated MEK signaling in Müller glia using *Rax-CreER^T2^;R26^LSL-MEK1DD^* mice. Remarkably, EdU-positive Müller glia were detected two weeks after MEK1DD induction, even in the absence of retinal injury (Fig. S3F). However, the overall number of proliferating cells remained low, indicating that sustained ERK activation alone is insufficient to drive robust regeneration. These observations suggest that ERK signaling functions as a critical potentiating mechanism that primes Müller glia toward a regenerative state, while additional injury-induced cytokine signals are required to fully activate proliferative responses.

### STAT Inhibition Enhances Regenerative Potential but Fails to Rescue Müller Glial Potentiation in the Absence of FGF Signaling

To investigate the mechanisms underlying the impaired regenerative response observed in FGFR-deficient Müller glia, we examined the activity of downstream signaling pathways. In addition to the expected reduction in ERK1/2 activation, conditional ablation of FGF signaling resulted in a marked increase in STAT3 activation in response to NMDA treatment of the retina (Fig. S4A-B”). This observation raised the possibility that FGF signaling promotes Müller glial regeneration, at least in part, by suppressing STAT activity.

To test this hypothesis, we examined whether pharmacological inhibition of STAT3/5 using SH-4-54 could stimulate Müller glial proliferation or enhance the activity of established regenerative stimuli. Interestingly, administration of SH-4-54 alone was sufficient to induce Müller glial proliferation, as evidenced by the presence of EdU+/tdTomato+ cells in tamoxifen-induced *Rax-CreER^T2^;Ai9* mice (Fig. 5A,B). These findings suggest that endogenous STAT signaling acts as a negative regulator of Müller glial cell-cycle re-entry. We next assessed whether STAT inhibition could synergize with other regenerative pathways. Notably, co-administration of SH-4-54 and Activin-A resulted in a dramatic increase in Müller glial proliferation compared with either treatment alone (Fig. 5B). This combination generated the most robust proliferative response observed throughout the study, indicating a strong cooperative interaction between Activin and STAT signaling pathways during Müller glial reprogramming.

**Figure 5.**
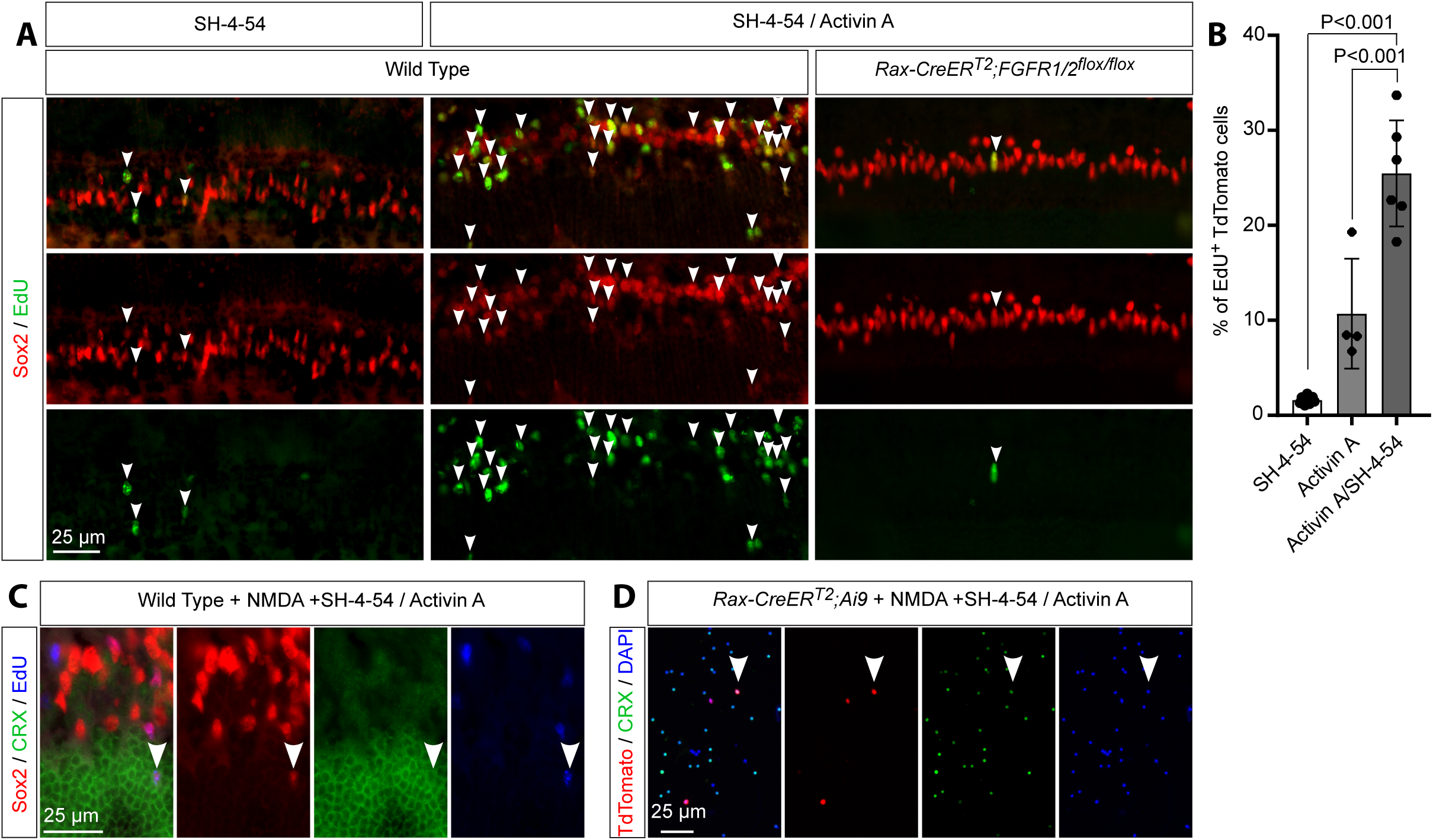
Combined STAT3/5 inhibition and Activin-A robustly promotes Müller glial regeneration. **(A)** Representative retinal flat mounts showing EdU incorporation following SH-4-54 alone or SH-4-54 plus Activin-A treatment after NMDA injury. **(B)** Quantification of EdU-positive tdTomato-positive Müller glia (mean ± S.D., One-way ANOVA, *n* = 3). **(C)** Immunostaining for Crx, Sox2 and EdU demonstrating acquisition of neurogenic marker expression following combined treatment. **(D)** Validation of Crx expression in lineage-traced Müller glia using dissociated *Rax-creER^T2^;Ai9* retinal cells.

To determine whether the proliferating Müller glia were acquiring neurogenic characteristics, we examined expression of the photoreceptor-associated transcription factor Crx. Immunohistochemical analysis revealed that a subset of EdU+/Sox2+ cells expressed Crx following combined SH-4-54 and Activin-A treatment (Fig. 5C). To further validate this finding, retinas from tamoxifen-induced *Rax-CreER^T2^;Ai9* mice receiving the combination treatment were dissociated into single-cell suspensions and immunostained for Crx. Consistent with the histological analyses, a proportion of tdTomato-positive Müller glia expressed Crx (Fig. 5D), indicating that these cells had initiated a neurogenic transcriptional program. Together, these data demonstrate that combined STAT3/5 inhibition and Activin-A stimulation not only promotes robust Müller glial proliferation but also enhances their regenerative potential.

Given the potency of this treatment paradigm, we next asked whether it could rescue the regenerative defect caused by loss of FGF signaling. However, despite generating a strong proliferative response in wild-type retinas, combined SH-4-54 and Activin-A treatment produced only a small number of EdU+/Sox2+ Müller glia in FGFR-deficient mice (Fig. 5A,B), indicating that suppression of STAT3/5 signaling is insufficient to fully restore regeneration in the absence of FGF signaling. Collectively, these findings identify STAT signaling as an important negative regulator of Müller glial proliferation and demonstrate that its inhibition can substantially enhance regenerative responses. However, the inability of STAT3/5 inhibition to rescue regeneration in FGFR-deficient Müller glia indicates that FGF signaling potentiates Müller glial regenerative competence through additional mechanisms beyond the regulation of STAT activity alone.

### Single-Cell Transcriptomic Analysis Reveals FGF-Dependent Regulatory Networks that Potentiate Müller Glial Regeneration

Given the robust proliferative response elicited by combined Activin-A and STAT3/5 inhibition, together with the inability of FGFR-deficient Müller glia to regenerate, we sought to identify the transcriptional mechanisms through which FGF signaling potentiates Müller glial regeneration. To this end, we performed single-cell RNA sequencing (scRNA-seq) on retinas harvested following SH-4-54 and Activin-A treatment (Fig. 6A). To preserve native transcriptional states and minimize artefacts associated with fluorescence-activated cell sorting, retinal cells were sequenced immediately following tissue dissociation without prior enrichment. A total of 30,088 cells from wild-type retinas and 41,445 cells from conditional FGFR knockout retinas were profiled, with an average sequencing depth of 879 genes per cell. All major retinal cell populations, including rods, cones, bipolar cells, horizontal cells, amacrine cells, and Müller glia, were identified (Fig. 6B). As expected, retinal ganglion cells were largely absent due to their depletion following NMDA-induced injury.

**Figure 6.**
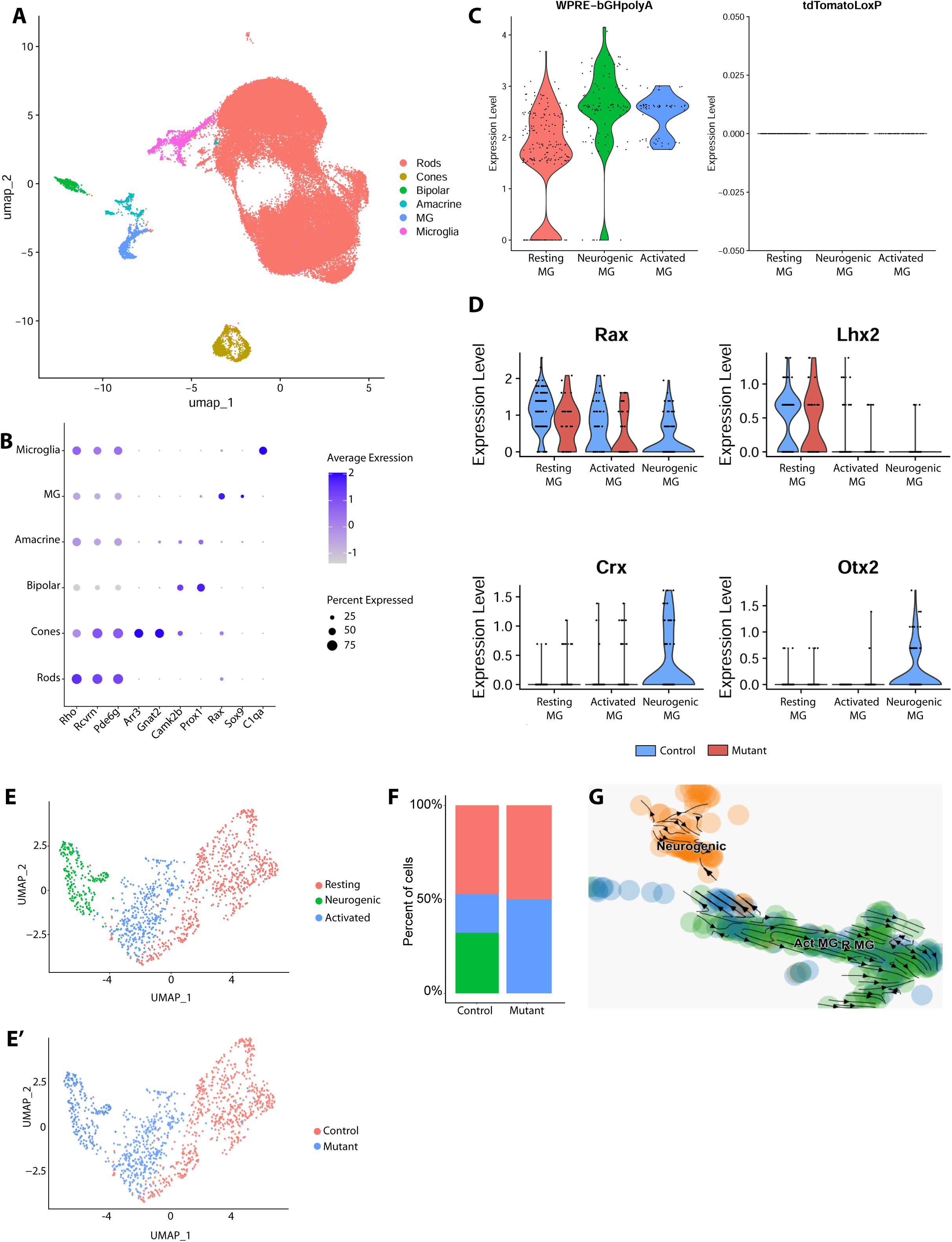
Single-cell transcriptomic analysis reveals impaired neurogenic reprogramming following FGF receptor deletion. **(A)** UMAP visualization of scRNA-seq data from control and *FGFR1/2* conditional knockout retinas following SH-4-54/Activin-A treatment. **(B)** Identification of retinal cell populations using established marker genes. **(C)** Isolation of lineage-traced Müller glia based on expression of the recombined Ai9 reporter transcript (WPRE-bGHpolyA) and exclusion of cells expressing the unrecombined tdTomato allele. **(D)** Differentially expressed genes defining resting, activated and neurogenic Müller glial populations. **(E)** Integrated UMAP of isolated Müller glia from both genotypes. **(F)** Relative abundance of Müller glial subpopulations. **(G)** RNA velocity analysis showing transitions from activated Müller glia toward either resting or neurogenic states.

To specifically analyse Müller glia and their progeny, we exploited the Ai9 lineage-tracing system to distinguish cells expressing the recombined tdTomato reporter transcript from those retaining the unrecombined locus (Fig. 6C). Cells expressing the recombinant WPRE-bGHpolyA transcript above a stringent threshold (>1.5 expression units) and lacking detectable expression of the unrecombined allele were retained for downstream analyses. These stringent filtering criteria were applied to minimize contamination arising from doublets or adherent cellular debris. Following filtering, 427 lineage-traced cells from wild-type retinas and 417 cells from conditional knockout retinas were recovered and integrated using the Seurat workflow. As anticipated, the majority of lineage-traced cells clustered within resting and activated Müller glial populations characterized by high Rax and high or low Lhx2 expression (Fig. 6D) ^53^. Consistent with the prior finding that tdTomato-positive Müller glia expressed Crx (Fig. 6D), an additional neurogenic Müller glial cluster was identified exclusively in wild-type retinas based on the expression of *Crx* and *Otx2*. Resting Müller glia represented approximately half of all lineage-traced cells in both conditions (Fig. 6E,F). In conditional knockout retinas, the remaining cells were almost exclusively classified as activated Müller glia. In contrast, wild-type retinas contained roughly equal proportions of activated and neurogenic Müller glia, indicating successful progression toward a regenerative state. RNA velocity analysis further revealed that while most activated Müller glia reverted toward a resting state, a subset underwent a transcriptional transition toward neurogenic differentiation (Fig. 6G).

Because this neurogenic transition was impaired in the conditional knockout sample, we further performed SCENIC analysis to identify the regulons that are responsible for this process. Interestingly, both activated and resting mutant Müller glia exhibited upregulation of the Hes1 regulon, which was not observed among the top regulons in wild-type Müller glia (Fig. 7A). We further investigated the differential gene expression of the activated and resting Müller glia between the mutant and control sample to identify the potential genes responsible for Müller glia potentiation. Consistent with expectations, *Hes1* was upregulated in mutant Müller glia, along with additional genes implicated in the inhibition of retinal regeneration, including *S1pr1, Cdkn1b, Rb1, Ep300* and *Crebbp* (Fig. 7B) ^54–56^. Among the identified candidates, S1pr1 was of particular interest because recent studies in zebrafish demonstrated that S1pr1 activation inhibits spontaneous retinal regeneration following injury ^57^. We therefore investigated whether aberrant S1pr1 signaling contributed to the regenerative failure of FGFR-deficient Müller glia. Wild-type retinas were treated with either the S1pr1 agonist SEW2871 in combination with SH-4-54 and Activin-A. Consistent with previous findings, activation of S1pr1 signaling significantly suppressed Müller glial proliferation (Fig. 7C,D). These results indicate that elevated S1pr1 signaling may contribute to the reduced regenerative competence observed following FGF depletion. Overall, the scRNA-seq analysis suggests that FGF signaling coordinately suppresses multiple anti-regenerative pathways, including *S1pr1, Cdkn1b, Rb1, Ep300* and *Crebbp*, thereby establishing a transcriptional landscape permissive for regeneration.

**Figure 7.**
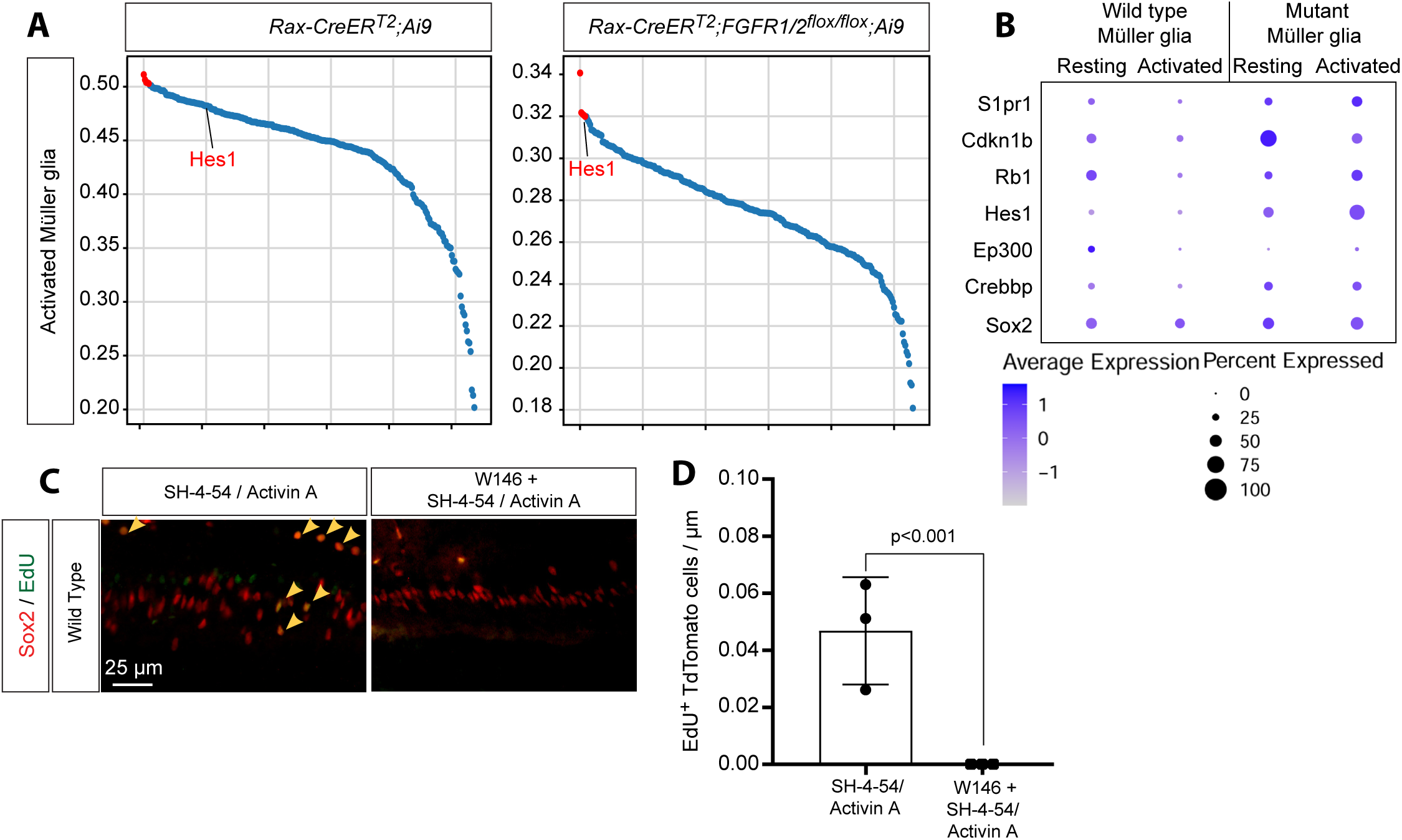
FGF signalling suppresses anti-regenerative transcriptional programs during Müller glial potentiation. **(A)** SCENIC regulon analysis comparing resting and activated Müller glia from control and *FGFR1/2* conditional knockout retinas. **(B)** Differential expression of genes associated with inhibition of retinal regeneration in *FGFR*-deficient Müller glia. **(C)** Representative retinal sections showing Müller glial proliferation following modulation of S1PR1 signalling. **(D)** Quantification of proliferating Müller glia shown as Sox2+/EdU+ cells (mean ± S.D., Two - sided Student’s t test, ns: not significant, *n* = 3).

## Discussion

In this study, we sought to resolve the longstanding uncertainty surrounding the role of FGF signaling during mammalian retinal regeneration. Our findings demonstrate that although FGF signaling alone is insufficient to induce Müller glial regeneration, it is nevertheless indispensable for this process. Specifically, neither exogenous administration of bFGF nor genetic overexpression of FGF8 was capable of inducing Müller glial proliferation in vivo, despite robust activation of the downstream ERK/MAPK pathway. These observations contrast with previous in vitro studies and findings from regenerative species such as zebrafish and chick, where FGF signaling has been reported to stimulate Müller glial proliferation and regeneration. While one potential limitation of intravitreal growth factor delivery is incomplete tissue penetration, our biochemical and histological analyses demonstrated robust ERK activation throughout the Müller glial population. Moreover, this response was abolished in FGFR1/2-deficient Müller glia, confirming both the effectiveness of the treatment and the specificity of the conditional knockout model. The discrepancy between our in vivo findings and previous in vitro studies may therefore reflect fundamental differences in the transcriptional and signaling states of cultured Müller glia compared with their native retinal environment, resulting in an exaggerated responsiveness to FGF under culture conditions.

A key finding of this study is that loss of FGF signaling abolished Müller glial proliferation irrespective of the regenerative stimulus employed. Activation of EGF, Wnt, Activin, or ROCK-associated pathways all failed to induce regeneration in FGFR-deficient Müller glia, indicating that FGF signaling functions as a central permissive regulator of regenerative competence rather than as a conventional mitogenic signal. These findings support a model in which FGF signaling primes Müller glia to respond to injury-induced regenerative cues. Similar observations have been reported in the chick retina, where depletion of microglia impaired Müller glial regeneration, yet exogenous FGF administration restored proliferative capacity ^47^. Interestingly, our single-cell transcriptomic analyses detected only limited FGF expression within microglia following injury. This raises the possibility that microglia either transiently express FGF during the acute injury response or alternatively regulate regeneration through upstream signals that stimulate FGF production by Müller glia themselves. Further studies will be required to define the cellular source of injury-induced FGF signaling within the retina.

Our mechanistic analyses further revealed that the regenerative function of FGF signaling is mediated predominantly through sustained activation of the ERK/MAPK pathway. Although multiple cytokines and growth factors can transiently activate ERK following retinal injury, inhibition of MEK/ERK completely abolished Müller glial proliferation, whereas inhibition of PI3K/AKT produced only a partial reduction in regeneration. Conversely, constitutive activation of MEK signaling was sufficient to induce limited Müller glial proliferation even in the absence of injury. Together, these findings suggest that prolonged rather than transient ERK activation is required to establish a regenerative state. We therefore propose that one of the primary functions of FGF signaling during regeneration is to maintain sustained ERK activity following injury, thereby preserving Müller glial competence to respond to additional pro-regenerative signals. This mechanism may also explain why exogenous FGF can compensate for the loss of microglia-derived signals in regenerative species.

Another important observation was the marked increase in STAT3 activation following FGF receptor deletion. STAT3 signaling has previously been implicated as a barrier to mammalian Müller glial regeneration, and our findings are consistent with studies demonstrating enhanced regenerative responses following STAT inhibition ^58^. Indeed, pharmacological inhibition of STAT3/5 was sufficient to induce Müller glial proliferation, and its combination with Activin-A generated the most robust regenerative response observed in this study. Notably, a subset of proliferating Müller glia induced by this treatment expressed the neurogenic transcription factor Crx, indicating progression toward a regenerative transcriptional state. However, despite the potency of this combination, STAT3/5 inhibition failed to fully rescue regeneration in FGFR-deficient Müller glia. These results indicate that suppression of STAT signaling represents only one component of the broader regenerative program regulated by FGF and suggest that additional FGF-dependent pathways are required to establish full regenerative competence.

To identify these pathways, we performed single-cell transcriptomic profiling of wild-type and FGFR-deficient Müller glia following regenerative stimulation. Interestingly, resting and activated Müller glial populations from both genotypes integrated closely following computational analysis, suggesting that the transcriptional consequences of FGF loss are relatively subtle. Nevertheless, the neurogenic Müller glial population identified in wild-type retinas was almost completely absent in FGFR-deficient samples, indicating a failure to progress toward a regenerative state. Differential expression and regulon analyses identified several candidate pathways that may underlie this defect. Among these, increased Hes1, S1pr1, p27, NF-κB, p300/CBP, and NFI activity were prominent features of FGFR-deficient Müller glia, all of which have previously been implicated in restricting retinal regeneration. Conversely, wild-type Müller glia displayed enhanced activity of FGF-responsive transcriptional programs, including Etv3, and showed reduced activation of multiple anti-regenerative networks.

One particularly intriguing observation was the reciprocal regulation of pathways associated with NF-κB activity. FGFR-deficient Müller glia exhibited increased expression of Nfkb1 and Nfkb2, whereas wild-type Müller glia displayed transcriptional signatures suggestive of reduced NF-κB signaling. This finding is notable because NF-κB activation has been repeatedly associated with reactive gliosis and suppression of Müller glial proliferation ^59^. Previous studies have demonstrated that STAT3 can prolong NF-κB activity through p300-mediated acetylation ^60^, while histone deacetylases such as HDAC6 can suppress NF-κB signaling by promoting its deacetylation and limiting nuclear retention ^61,62^. Based on these observations, we propose a model in which sustained FGF signaling potentiates Müller glia by simultaneously suppressing STAT3, p300/CBP, Hes1, and NF-κB activity while promoting regenerative transcriptional programs downstream of ERK signaling. Through this coordinated regulation, FGF signaling shifts Müller glia away from a gliotic state and toward a regeneration-competent state capable of responding to injury-induced cues.

## Acknowledgements

The authors thank Drs. Seth Blackshaw, Liang Ma and David Ornitz for mice. The work was supported by grants from NIH (EY017061 and EY018868 to X.Z.). N.M. is supported by a Knights Templar Eye Foundation Career Starter Grant Award. The Columbia Ophthalmology Core Facility is supported by NIH Core grant 5P30EY019007 and unrestricted funds from Research to Prevent Blindness (RPB).

## Tables

**Table S1.**
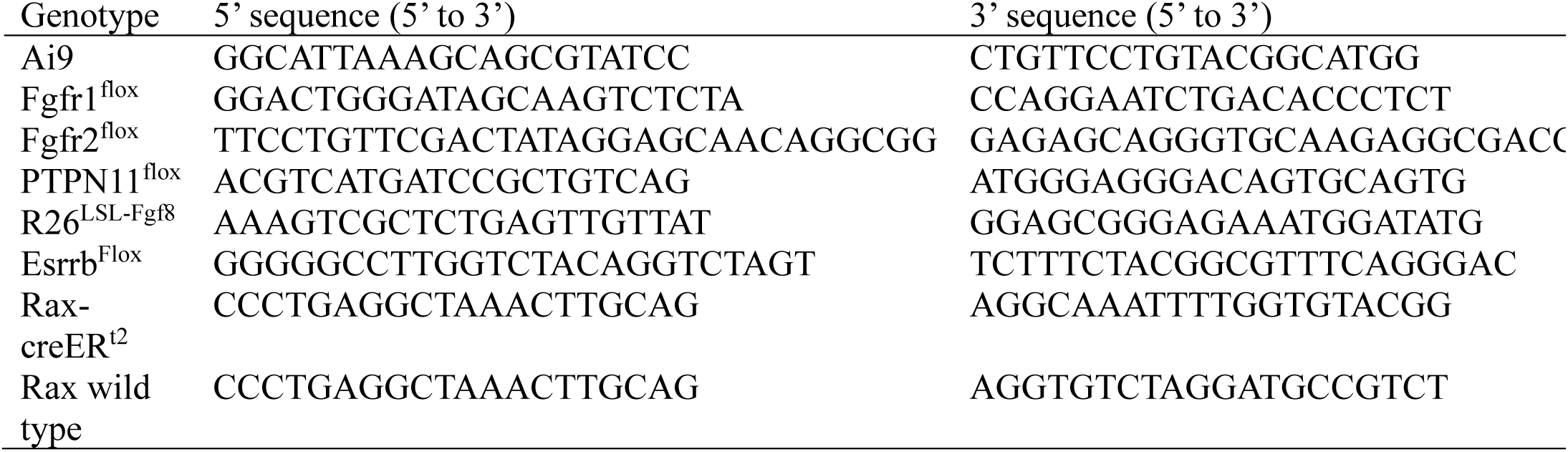
Genotyping oligos.

**Table S2.** Chemical compounds used for intraocular injection.

| Compound | Concentration | Company | CatLog no. |
| --- | --- | --- | --- |
| Activin A | 2 µg/µl | R&D Systems | 338-AC |
| Chir99021 | 0.03 µM | Tocris Bioscience | 252917-06-09 |
| EGF | 1 µg/µl | R&D Systems | 2028-EG |
| Diethylstilbestrol | 10 µM | Sigma-Aldrich | D4628 |
| DY131 | 10 µM | MedChemExpress | HY-15483 |
| bFGF | 1 µg/µl | ScienCell | 104-02 |
| LDN193189 | 0.2 µM | ReproCell | 04-1974 |
| LY294002 | 20 µM | Tocris Bioscience | 1130 |
| NMDA | 100 mM | Sigma-Aldrich | 6384-92-5 |
| PD0325901 | 10 µM | Tocris Bioscience | 391210-10-9 |
| SB431542 | 0.1 µM | Santa Cruz | sc-204265A |
| SB525334 | 0.1 µM | Selleckchem | S1476 |
| SEW2871 | 0.1 µM | MedChemExpress | HY-W008947 |
| TGFb1 | 1 µg/µl | Cell Signaling | 5231LC |
| W146 | 0.1 µM | MedChemExpress | HY-101395 |

**Table S3.** Antibodies used in the study.

| Antibody | Species | Dilution factor | Method | Company | CatLog no. |
| --- | --- | --- | --- | --- | --- |
| AKT | Mouse | 1:800 | WB | Cell Signaling | 2920 |
| Brn3a | Mouse | 1:200 | IHC | Chemicon | MAB1585 |
| CRX | Rabbit | 1:500 | IHC | Atlas Antibodies | HPA036762 |
| Cyclin D3 | Mouse | 1:200 | IHC | Cell Signaling | DCS22 |
| ERK1/2 | Rabbit | 1:2000 | WB | Cell Signaling | 4695 |
| Esrrb | Rabbit | 1:100 | IHC | Abcam | ab150539 |
| GFAP | Rabbit | 1:1000 | IHC | Dako | Z0334 |
| IBA-1 | Rabbit | 1:1000 | IHC | Wako Chemicals | 019-19741 |
| Lef1 | Rabbit | 1:200 | IHC | Cell Signaling | 2230 |
| Otx2 | Goat | 1:500 | IHC | R&D systems | AF1979 |
| p-p44/42<br>MAPK | Rabbit | 1:100 | IHC | Cell Signaling | 4370 |
| p-p44/42<br>MAPK | Mouse | 1:1000 | WB | Santa Cruz | SC-7383 |
| p-AKT | Rabbit | 1:100 &<br>1:1000 | IHC &<br>WB | Cell Signaling | 9271 |
| p-Histone H3 | Rabbit | 1:200 | IHC | Cell Signaling | 9701 |
| p-Stat3 | Rabbit | 1:1000 | WB | Cell Signaling | 9134 |
| p-PLCγ1 | Rabbit | 1:1000 | WB | Cell Signaling | 14008 |
| p-Smad2 | Rabbit | 1:1000 | WB | Abcam | ab53100 |
| PLCγ1 | Mouse | 1:1000 | WB | BD Transduction<br>Laboratory | 610028 |
| Sox2 | Rat | 1:400 | IHC | Thermo Fisher | 14-9811-82 |
| Sox9 | Rabbit | 1:200 | IHC | Cell Signaling | 82630 |
| Stat3 | Mouse | 1:1000 | WB | Cell Signaling | 9139 |

## Figure Legends

**Figure S1.**
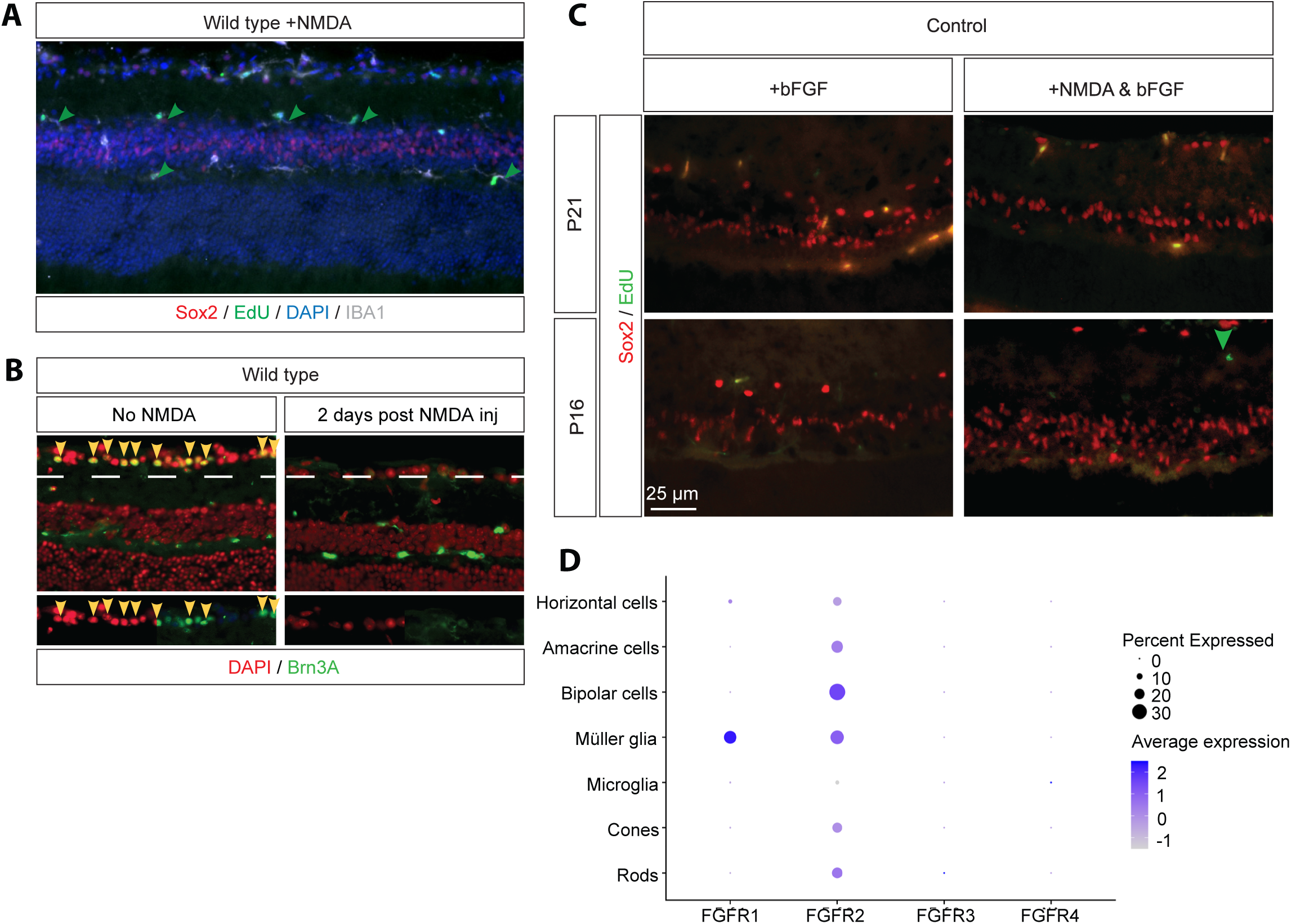
Characterization of retinal regeneration after NMDA injury and bFGF treatment. **(A)** EdU incorporation following NMDA injury identifies proliferating IBA1-positive microglia rather than Sox2-positive Müller glia. **(B)** NMDA-induced depletion of retinal ganglion cells visualized by loss of Brn3a staining. **(C)** Assessment of Müller glial proliferation in P16 and P21 mice following bFGF treatment with or without NMDA injury. **(D)** Dot plot showing expression of FGF receptor genes in the adult mouse retina.

**Figure S2.**
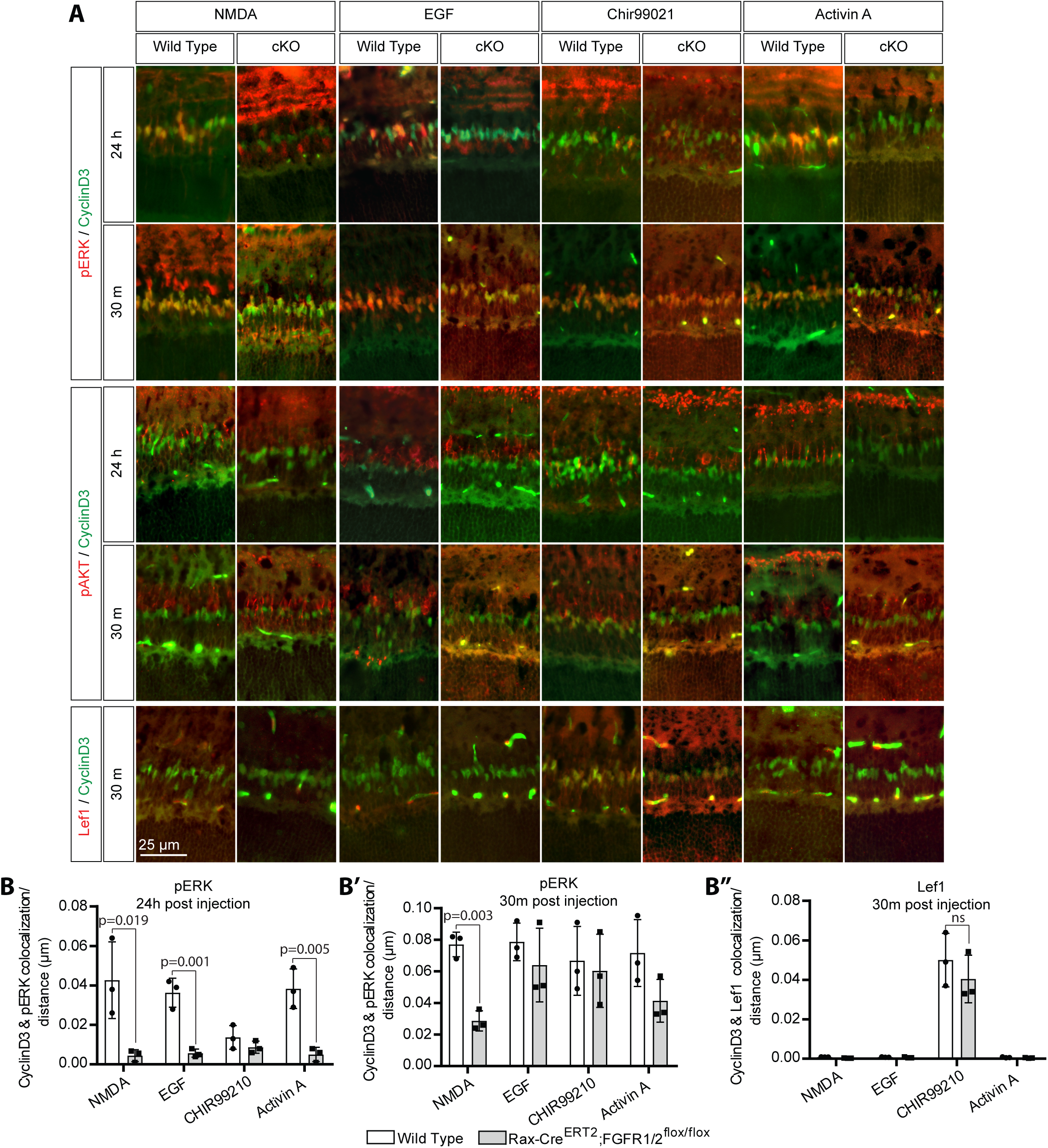
Sustained ERK activation downstream of FGF signalling. **(A)** Immunostaining for pERK, pAKT and Lef1 in control and FGFR1/2 conditional knockout retinas following NMDA, EGF, CHIR99021 or Activin-A treatment at 30 min and 24 h. **(B)** Quantification of CyclinD3 colocalization with pERK, pAKT or Lef1 (mean ± S.D., One-way ANOVA, ns-non significance, *n* = 3).

**Figure S3.**
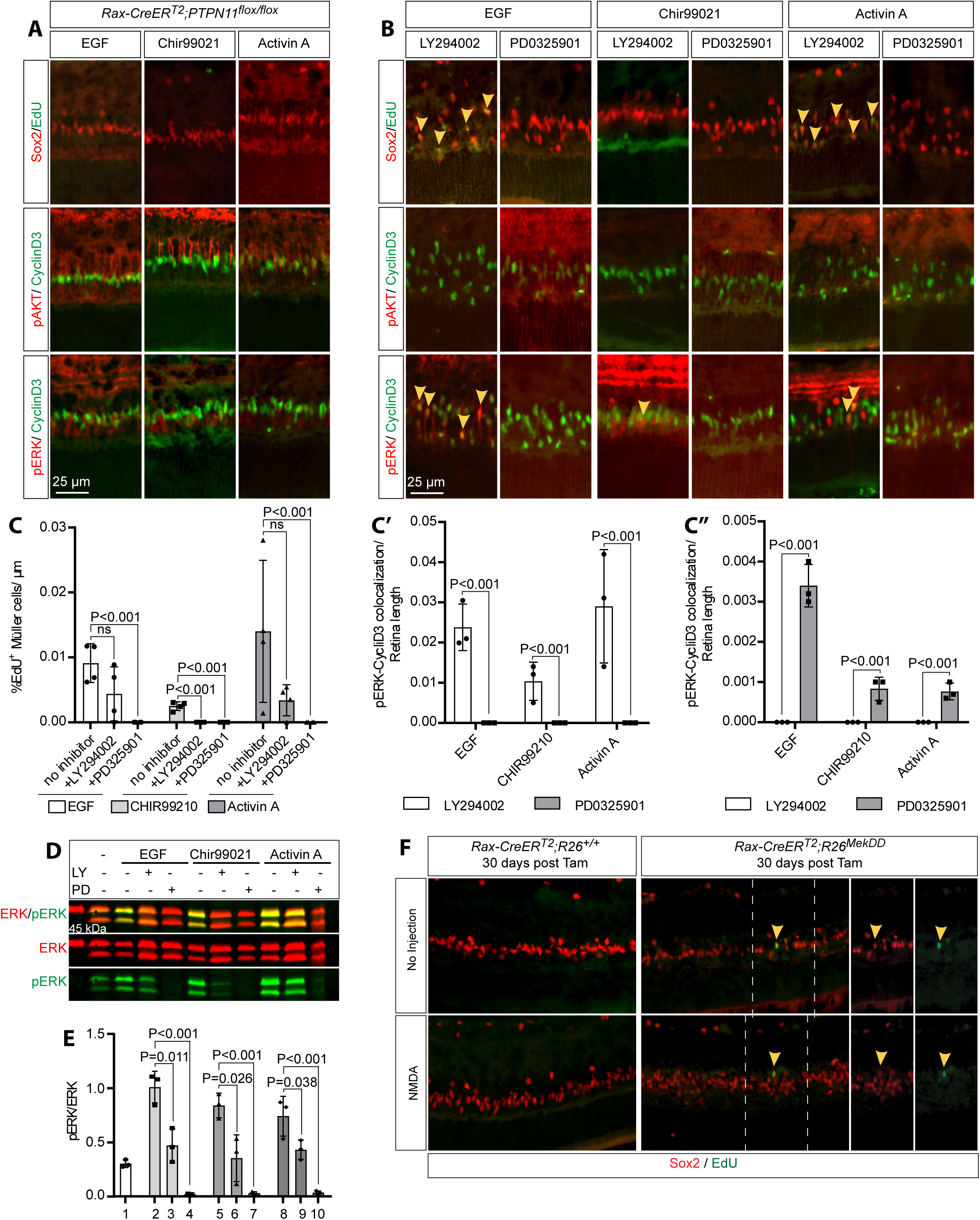
Müller glial regeneration requires MEK/ERK signalling. **(A)** EdU incorporation and pERK/pAKT immunostaining in control and Shp2 conditional knockout retinas following EGF, CHIR99021 or Activin-A treatment. **(B)** Effects of LY294002 or PD0325901 on regeneration induced by EGF, CHIR99021 or Activin-A. **(C)** Quantification of Sox2+/EdU+, CyclinD3+/pERK and CyclinD3+/pAKT cells (mean ± S.D., One-way ANOVA, *n* = 3). **(D)** Representative pERK immunoblot from retinas treated with regenerative stimuli in the presence of pathway inhibitors. **(E)** Quantification of pERK relative to total ERK (mean ± S.D., One-way ANOVA, *n* = 3). **(F)** EdU and Sox2 staining of *Rax-creER^T2^*;*R26^LSL-MEK1DD^*retinas following MEK1DD activation.

**Figure S4.**
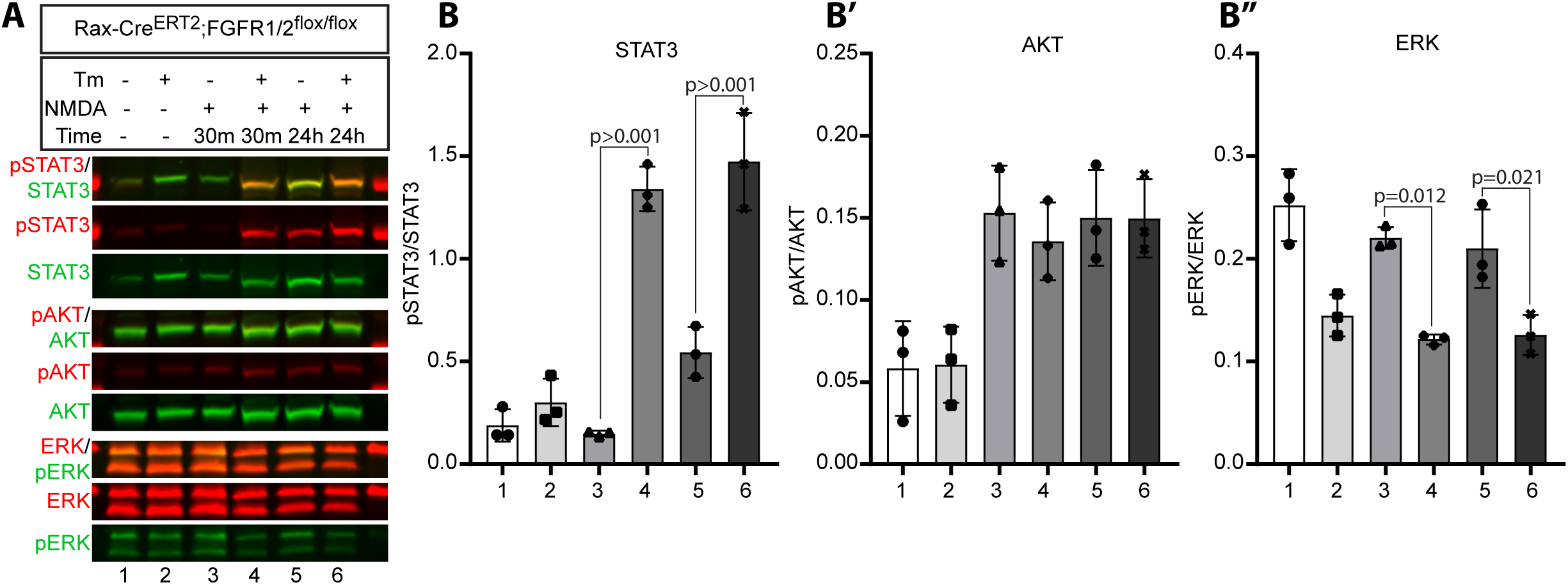
STAT signalling is enhanced following loss of FGF signalling. **(A)** Representative immunoblots of pSTAT3, pERK and pAKT in retinal lysates from control and *FGFR1/2* conditional knockout mice collected 30 min or 24 h after NMDA injury. **(B)** Quantification of phosphorylated protein levels relative to total protein (mean ± S.D., One-way ANOVA, *n* = 3).

